# Modeling the mosaic structure of bacterial genomes to infer their evolutionary history

**DOI:** 10.1101/2023.09.22.558938

**Authors:** Michael Sheinman, Peter F. Arndt, Florian Massip

## Abstract

The timing and phylogeny of bacterial evolution is difficult to reconstruct because of a scarce fossil record, deep genomic divergences and complexities associated with molecular clocks. Studying bacterial evolutionary history using rich and rapidly accumulating genomic data requires accurate modeling of genome evolution, taking into account that different parts of bacterial genomes have different history. In particular, along the genome, different loci are subject to different selective pressure. In addition, some are horizontally transferred from one bacterium to another, resulting in a mosaic-like genome structure. An important technical aspect is that loci with high effective mutation rates can diverge beyond the aligner detection limit, biasing the genome-wide divergence estimate towards more conserved loci. Therefore, the genome-wide molecular clock cannot be directly applied to study bacterial evolutionary history. In this article, we propose a novel method to gain insight into bacterial evolution based on statistical properties of genomic sequences comparisons. The length distribution of the sequence matches is shaped by the effective mutation rates of different loci, by the horizontal transfers and by the aligner sensitivity. Based on these inputs we build a model and demonstrate that it accounts for the empirically observed distributions, taking the *Enterobacteriaceae* family as an example. Using the model and the empirical data we fit the evolutionary parameters: time divergences and horizontal transfer rates. Based on the estimated time divergences we build a time-calibrated phylogenetic tree, demonstrating the accuracy of the method and its ability to unravel vertical and horizontal transfers in bacterial genomes.

## I. Introduction

Reconstructing bacterial evolution is a challenging task. In contrast to multicellular organisms for which an abundant fossil record helps to date events on phylogenetic trees, bacteria leave very little trace of their existence [1]. Despite the accumulation of genomic data in the last decades, divergence times of many bacterial taxa are yet to be reliably estimated. Such estimates may be very useful, especially when combined with host, habitat or ecosystem data [2]. The concept of the “molecular clock” [3, 4] has become fundamental in the analysis of evolution [5, 6], but applying it to date bacterial diversification events is often problematic. In particular, it is necessary to determine the rate at which nucleotides mutate over time, *i*.*e*. the speed at which the clock “ticks”. However, this effective mutation rate does not only depend on the background point mutation rate (associated with replication errors and repair) and on the generation time of the bacterium [7, 8], but also on different ecological parameters [9, 10], location along the chromosome [11–13], activity of nucleoid-associated proteins [14], fitness effects of the mutations [15–17] and other factors. All this, being difficult to assess in practice, prevent an accurate estimation of divergence times. Furthermore, the molecular clock is also obfuscated by horizontal gene transfers, especially if the clock is based on a small number of genes (*e*.*g*. slow evolving rRNA genes) [18–22] and some of them have taken part in horizontal transfer [23–27]. Finally, loci with high effective mutation rates diverge rapidly, such that alignment algorithms do not detect such homologous loci in distant bacteria. As a consequence, these regions are not considered in divergence time estimation, resulting in information loss and underestimation of the divergence time.

While the phylogenetic relationships between species can usually be inferred using the molecular clock, we lack reliable and scalable methods to infer the branching times on phylogenetic trees. Namely, the bacterial molecular clock cannot be satisfactorily calibrated in contrast to multicellular organisms [28]: one has to relate to ecological events at known times to specific points in the phylogenetic tree [29]. For instance, to link the evolution of bacteria and their hosts [30], *etc*. [6]. However, this approach often leads to orders of magnitude discrepancies between estimates of the mutation rate on different timescales [6, 28, 31, 32]. Such discrepancies led to the hypothesis of time-dependent mutation rate [33–36] and corresponding relaxed molecular clock models [37]. Such relaxed molecular clock models however require to fit many free parameters, which can be problematic when there is only limited amount of data available. Moreover, these models are mostly applied to a small number of marker genes, such that large parts of the genomes are not considered in the time divergence estimations, discarding potentially useful information.

In sum, evolutionary reconstruction in bacteria is particularly difficult due to the mosaic structure of genomes: different loci evolve with different effective mutation rates, while some loci are acquired via horizontal gene/allele transfer. The mosaic structure of bacterial genomes can be directly observed. In the alignment of two bacterial genomes (see Figs. S1 and S2 for the example of *E. coli* and *S. enterica* pair) the mutation densities significantly vary along the alignment from one locus to another.

In this article, we show that taking into account the mosaic structure of genomes allows to resolve time estimate discrepancies and to reconstruct bacterial evolutionary history. We do so introducing the “mosaic molecular clock”, that is a molecular clock which accounts for variations of mutation rate from one locus to another. We show that quickly mutating loci cross the homology detection limit of the aligner sooner than loci mutating slowly and demonstrate how this affects empirical time divergence estimates. This model makes it possible to estimate time divergences based on all detectable homologous loci in the studied genomes, in contrast to most existing methods based on marker genes. Importantly, our model also takes into account horizontal gene transfer. Predictions of the model—total homologous region length, average similarity observed and other statistical properties—agree well with the empirical results for real taxa over a wide range of pairwise time divergences.

## II. Model and analytical solution

### The mosaic molecular clock model

The main assumption of our model is that bacterial genomes have a mosaic structure, that is, each locus *i* mutates with a different effective mutation rate *μ*_*i*_ and is inherited vertically. We refer to this part of the genome as *vertical*. By “Effective” mutation rate we mean the rate of mutation and its fixation in the population. In addition, a taxa pair horizontally exchanges random loci with rate *ρ*. The part of the genome that comprises such loci is denoted as *horizontal*. One can see a schematic illustration of the model in Fig. 1(a). Below we describe in more detail our assumptions about the effective mutation rate distribution and how we model detection limit of the aligner.

**FIG. 1:**
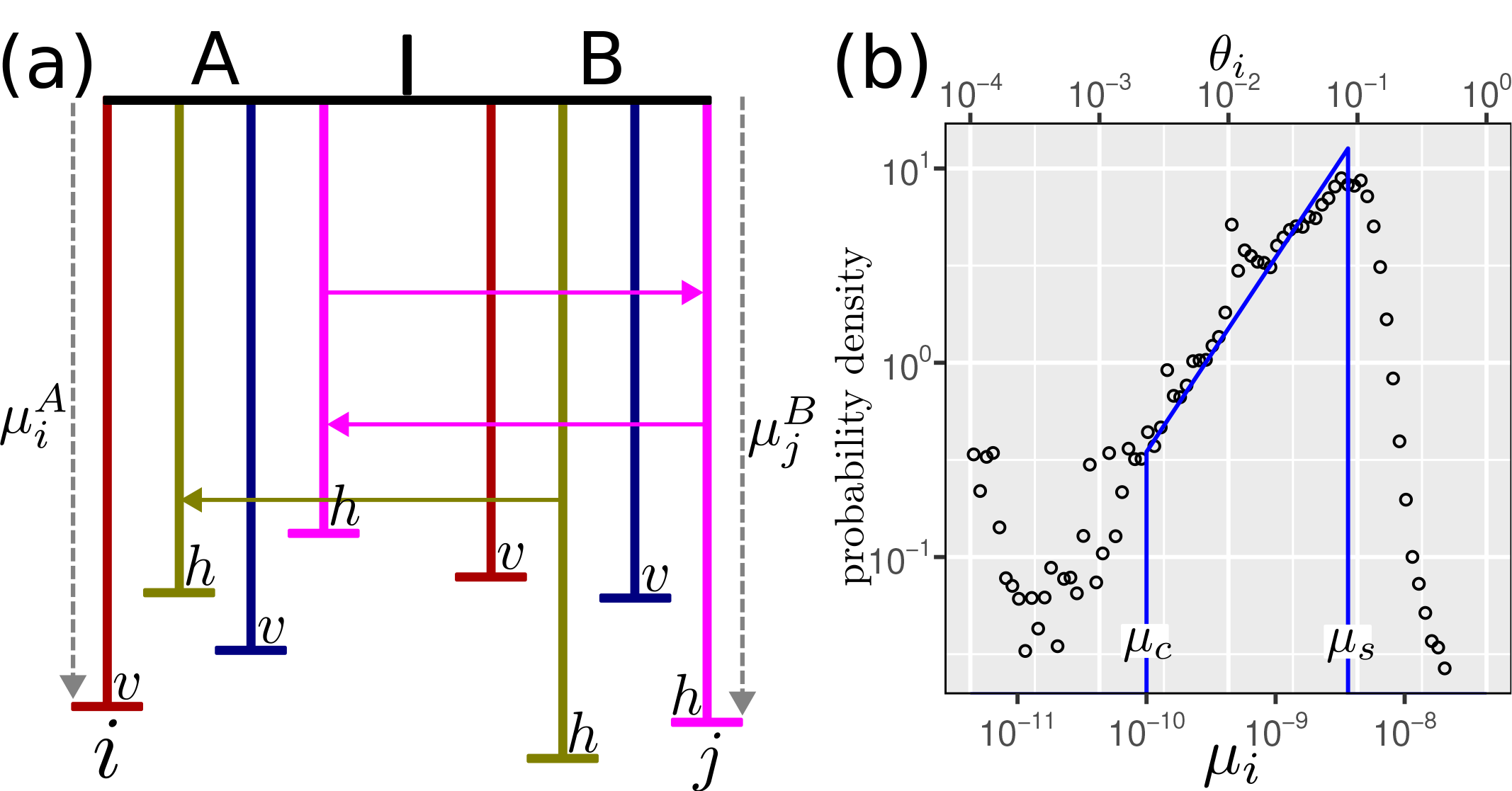
Illustration of the model for the evolution of two taxa, A and B. (a) Vertical lines represent different loci along the genome, evolving with different effective mutation rates, while horizontal lines depict horizontal transfer of loci between the taxa. Vertical and horizontal parts of the genomes are marked by *v* and *h*, correspondingly. (b) Distribution of mutation rate in a bacterial genome. Solid lines represent the assumed distribution of mutation rate, as described in Eq. (1) with *μ*_*c*_ = 10^*−*10^ [29] and *μ*_*s*_ = 3.64*·*10^*−*9^ [40]. Circles represent empirical distribution of segment divergences between *E. coli* and *E. albertii* (see upper horizontal axis). Segments were obtained using the `segmut R` package [41] (see Method). Here, to get the values of *μ*_*i*_ = *τθ*_*i*_ from the mutation densities *θ*_*i*_ we took *τ* = 2.3 *·* 10^7^, as obtained in further analyses (see Fig. 2(a)).

### Mutation rate distribution

The mutation rates are distributed between two values: the smallest one, *μ*_*c*_ corresponds to the mutation rate of the most *conserved* regions, like rRNA genes, while the largest one, *μ*_*s*_, corresponds to the *spontaneous* background point mutation rate. Relatively rare cases of faster than background mutation rate due to positive selection are ignored within the considered model. We model the mutation rate along the genome as a random variable. Its distribution is a crucial ingredient in the model that we infer using a combination of analytical arguments and empirical evidences.

Consider a locus *i* in two bacterial taxa A and B. The genomic divergence between the two bacteria along this locus is given by *θ*_*i*_ = *μ*_*i*_*τ* (in the *θ*_*i*_*≪*1 regime) where 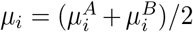 is the average of the two effective mutation rates, 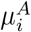 and 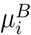, at this locus for these lineages and *τ* is the time divergence between the bacteria (twice the time to their last common ancestor). We assume that mutation rates of two bacterial lineages are not correlated (see discussion about this assumption in the Discussion and Summary section). Following this assumption, it can be demonstrated that the distribution of *μ*_*i*_ scales linearly for small values [38, 39]. Thus, it is expected that the divergences of different loci *θ*_*i*_ (in the vertical part of the genome) are also distributed in a similar linear fashion. We observe this linear regime empirically, as shown in Fig. 1(b).

In sum, our assumption for the distribution of the effective mutation rate is given by

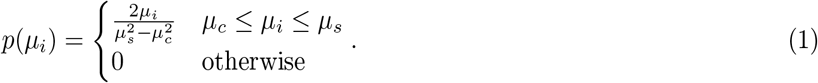

Further on, we omit the locus index *i* to simplify the notation.

### Alignment software detection limit

The ability of an alignment software to detect homology depends on the properties of the alignment algorithm used and on the properties of the considered sequences. Here we summarize the properties of the aligner into one effective parameter *δ*, assuming that an aligner can detect homologous sequences as long as their divergence *θ* is smaller than a threshold *δ*. Namely, if a given locus mutates with a mutation rate *μ* and the time divergence of this locus is *τ*, it is detected as homologous if and only if *μτ* ≤ *δ*. Defining *μ*_*a*_ as the mutation rate of the least conserved alignable region, for 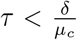 we have 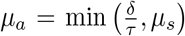. For 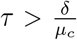 no homology of the vertical part of the genome can be detected. The dependence on the length is much weaker and is ignored.

We verified the validity of this simplifying assumption within our framework using simulated data and calculated the effective value of *δ*for the specific software that we used in this study (see discussion below in Section II A and Figs. S7,S8).

### A. Analytical solution

#### How to relate the model to genomic data?

Inferring the parameters of our model from empirical genomic data is challenging. This is due to the fact that, unlike artificial mosaics, the boundaries of the constituent pieces are not well recognizable and the pieces are often too small to be analysed thoroughly and even identified. Here we briefly discuss the validation procedure of our model using empirical data and demonstrate how to relate *p*(*μ*) to an easily accessible empirical quantity.

In principle, using the molecular clock assumption, the evolutionary time divergence *τ* between two DNA loci with effective mutation rate *μ* is related to the density of mismatches at these loci, *μτ* . To take into account the mosaic structure of the genomes, we consider a combination of clocks, one per locus—each clock ticking at a different pace due to the different mutation rates. In addition, some loci may also have undergone horizontal gene transfer, and thus vary in their divergence times. In this paper, we combine these different molecular clocks into one “mosaic molecular clock”.

In practice, the mosaic structure of the genomes we study is not known *a priori*, that is, one has to infer the regions with constant mutation densities. To this end, we developed a simple method called `segmut` to partition the genomes into regions with constant mismatch densities using a *χ*^2^ approach (see Methods and Figs. 1(b),S3,S2). This method has several drawbacks since it is computationally intensive, and the results are difficult to verify on empirical data. To circumvent this difficulty we decided to use another approach that was already efficiently applied in different contexts (see [38, 39, 42–44]). The main idea, we employ here, is to study the length distribution of maximal exact matches between homologous sequences. Indeed, one can show [45], using the result derived in [46] that studying match length distribution (MLD) allows to assess time divergence between DNA sequences. Namely, for a given *τ* and *μ* between two loci of length *K≫r* the expected number of their exact sequence matches, *m*(*r*), is given by

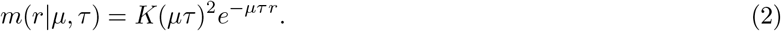

Therefore, if the mutation rate follows a certain distribution *p*(*μ*), the MLD for two genomes of length *L* is given by

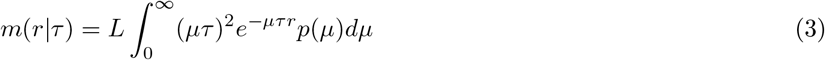

We note that the integral above can be represented as a Laplace transformation and:

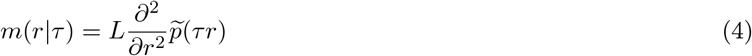

where 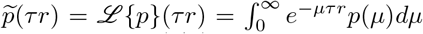 is the Laplace transform of *p*(*μ*). Hence, there is a direct relationship between the MLD *m*(*r*|*τ*) and the Laplace transform of the mutation rate distribution. As a consequence, studying *m*(*r*) – a quantity that can be easily computed for empirical data – allows to reconstruct the evolutionary history of the genomes, in particular their mutation rate distribution. Below we further take into account that *τ* is also distributed along genomes due to horizontal transfers, but the principle is the same: the distribution *m*(*r*) is easy to compute empirically, easy to calculate analytically and contains informations about the distributions of *μ* and *τ*.

To demonstrate the validity of our approach on empirical data, we computed the alignment of one strain of *E. coli vs*. one strain of *S. enterica* (Fig. S3(a)). Ignoring the mosaic structure of genomes, assuming that the average genome-wide density of mutations *θ* is uniform along the genome, the MLD would simply follow *m*(*r*) = *Lθ*^2^*e*^*−θr*^, which is very different from the empirical observation. To make sure that this discrepancy is due to the existence of loci with different effective mutation rates along the genome, we used the `segmut R` package (see methods) to reconstruct the mosaic structure of the genomes. This way we can compute the empirical distribution of mutation rates *p*(*θ*_*i*_). Assuming that the mutations are uniformly distributed inside each locus (that is, 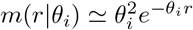) we can compute a pseudotheoretical MLD *m*(*r*) =Σ_*i*_ *p*(*θ*_*i*_)*m*(*r*|*θ*_*i*_) which very closely mimics the empirical distribution, demonstrating the validity of our model. The disagreement between the naive model (uniform distribution of mutations along the whole genome) and our mosaic model is even more evident when one considers the comparisons of many pairs of strains as shown for the analysis of all *vs*. all alignments of *E. coli vs. S. enterica* (Fig. S3(b)).

In the following, we use our model to calculate analytically the MLD, compare it to the empirical one and infer the model parameters *τ* and *ρ*for all considered pairs of taxa. To simplify the analysis, below we consider separately the MLD from the vertical part of the genome, *m*_*v*_, and the MLD from the horizontally transferred part, *m*_*h*_. In the next section, we calculate analytically the shape of the MLD for the vertical and horizontal parts of the genomes.

### Vertical part

The MLD from the *δ*-detectable vertical part of the genome (homologous loci with divergences smaller than *δ*) with time divergence *τ*, using Eq. (1), is given by

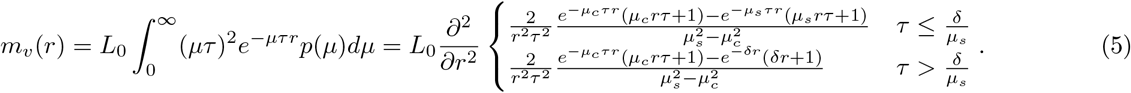

One can see that the tail of the MLD from the vertical part scales as *r*^*−*4^, as previously observed in eukaryotes [38, 39].

The total length of the *δ*-detectable homologous vertical part of the genome decreases with increasing time divergence*τ* and is given by

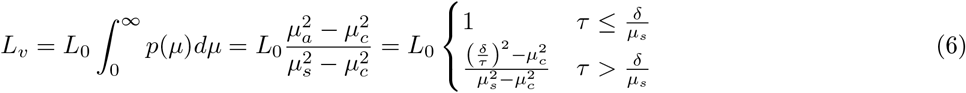

Along the vertical region with this length the average divergence is given by

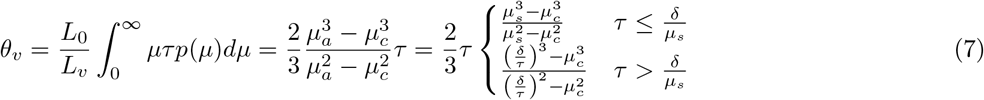

See Fig. S11 for representative plots of these functions.

### Horizontally transferred part

Assuming that only a small fraction of the genome has been transferred (*i*.*e*.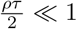), the MLD from the *δ*-detectable horizontal part of the genome can be written as:

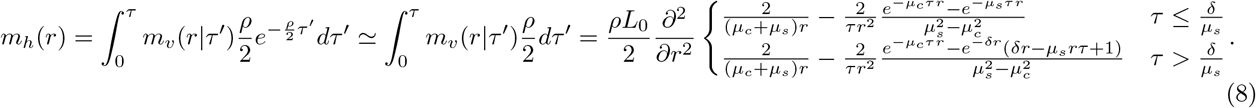

One can see that the tail of the MLD from the horizontally transferred part scales as *r*^*−*3^, as was also derived and shown empirically in Ref. [44].

In the same regime, the total length of the *δ*-detectable homologous part of the genome due to HGT is given by

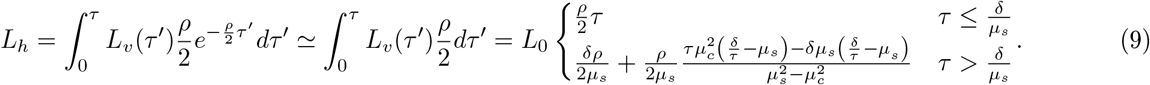

Along the horizontally transferred region with this length the average divergence is given by

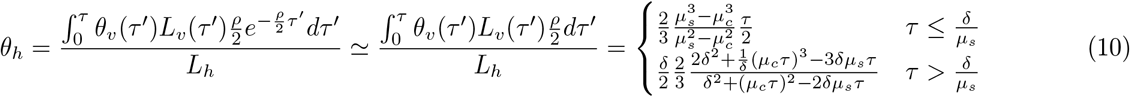

### Total MLD

Assuming that the total contribution of the homologous part due to HGT is much smaller than the contribution from the vertical part (*i*.*e. L*_*h*_*≪L*_*v*_), the total MLD is given by:

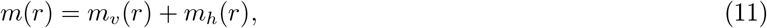

where *m*_*v*_ and *m*_*h*_ are given by Eqs. (5,8). The total *δ*-detectable homologous length is given by

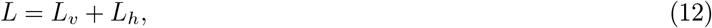

where *L*_*v*_ and *L*_*h*_ are given by Eqs. (6,9). The average divergence is given by

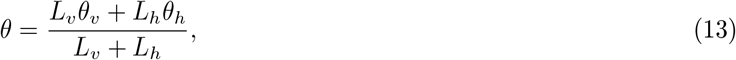

where *θ*_*v*_ and *θ*_*h*_ are given by Eqs. (7,10).

Finally, note that in our framework gene losses reduce the overall length of the homologous regions between the two taxa *L*_0_, but do not change the shape of the MLD.

### Numerical validation

To test our theory, we simulated the evolution of species under the assumptions of the model (see Fig. 1(a)): each locus mutates with a certain rate, distributed as (1) and horizontal transfer of loci occurs with rate *ρ*(see Methods). We aligned sequences obtained in these simulations using the `nucmer` software [47] with default parameters. Importantly, we kept only matches that are unique in both strains to reduce the influence of paralogs that are not the main focus of this study (see Methods). Fitting all MLDs with *δ*as a free parameter, we find that for the used aligner, *δ≃*0.25 results in good fits (see Fig. S7). Using this parameter we were able to fit the numerical MLDs, demonstrating the validity of our approach. In addition, the estimations of divergence times and horizontal transfer rates reproduced well the simulation parameters (see Fig. S8) showing that our method allow to reconstruct the evolutionary history of species from genomic data. In the following, analyzing empirical data we use *δ*= 0.25, assuming that *δ*is the property of the aligner and does not depend strongly on the analyzed sequences.

## III. Empirical validation

### MLD fitting—two regimes

We then tested our model on empirical data. We downloaded 3, 249 fully assembled genomes from 11 taxa of the *Enterobacteriaceae* family, as well as 759 genomes from two outgroups *Serratia* and *Vibrio* genera (see Table I for details).

We computed one MLD per pair of taxa from all 13×12/2 pairwise comparisons (see Methods for technical details, Fig. 2 for a few examples and Supplementary File mAll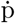df for all comparisons). Obtained MLDs exhibit two different regimes, corresponding to short and long matches, in good agreements with the prediction of our model. Indeed, we observe that short matches follow a power-law with a *−*4 exponent, as expected for the matches from the vertical part (see Eq. 5) while long matches are distributed according to a*−*3 power-law, as predicted for horizontally transferred segments (see Eq. 8). The location of the transition between the two regimes depends on the time divergence between the taxa: the closer the two taxa are, the longer the matches of the vertical part. Analytical prediction for the combination of the vertically and horizontally transferred part Eq. (11) fits well the empirical data for almost all taxa pairs.

**FIG. 2:**
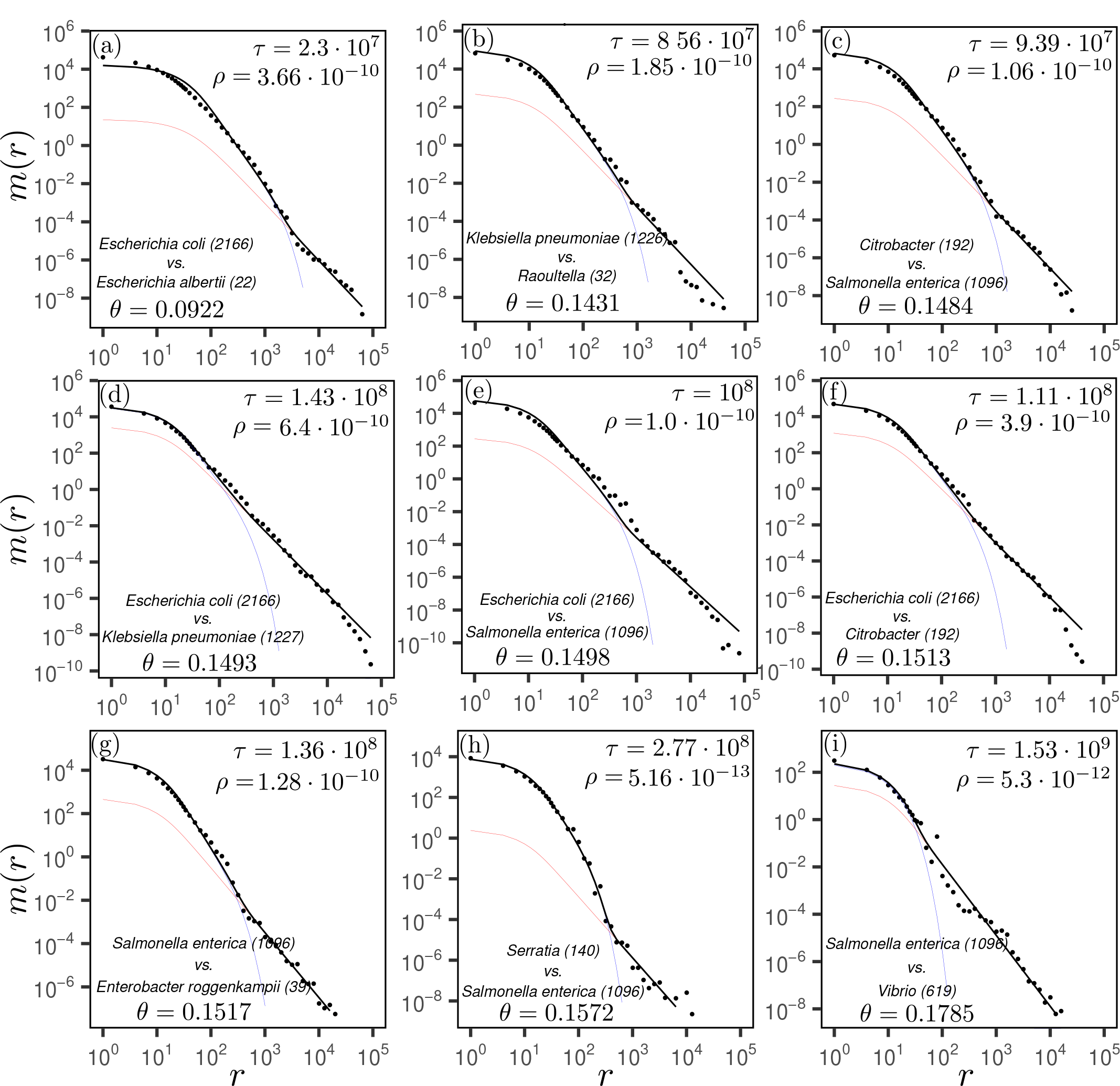
Match length distributions of 9 selected pairs of taxa. Names of the taxa are indicated in the bottom-left corner of the panels. The numbers in the brackets indicate number of strains. The average divergence is indicated by *θ*. The empirical data (dots) are fitted with Eq. (11) (black solid lines) using the global parameters *μ*_*s*_ = 3.64·10^*−*9^/bp/yrs, *μ*_*c*_ = 10^*−*10^/bp/yrs and *δ*= 0.25. The values of *τ* and *ρ*are fitted for each pair separately and are shown in the top-right corner. Using the obtained parameters the vertical and the horizontal parts of the match length distributions is plotted using Eqs. (5) (blue lines) and (8) (red lines), respectively. The genome length of the most recent common ancestor of two taxa is assumed to be the minimum of the taxas’ genome lengths.

### Model setbacks

Our model failed to estimate the time divergence for two specific taxa pairs: 1. *E. coli vs*.*E. fergusonii* and 2. *E. asburiae vs. E. hormaechei*. In Fig. S10 one can see the reason for this: the MLD of the first pair has a *m*(*r*)*∼r*^*−*3^ regime in the vertical part, implying that Eq. (1) is not valid and suggesting instead *p*(*μ*)*∼μ*^0^ for this pair. One can observe similar behaviours for another closely related pair in Fig. 2(a) (for this pair the time divergence estimate is nevertheless reasonable). For the other pair, *E. asburiae vs. E. hormaechei* (see Fig. S10(b)), the rate of horizontal transfer is that high that the horizontally transferred segments dominate over the vertical part. Since time divergence estimation is based solely on the vertical part, the signal is obfuscated. In fact, this demonstrates another quality of our approach: using MLD one can easily diagnose pairs for which the assumptions of the model are not fulfilled, and therefore the parameter estimation fails.

### Eukaryotic genomes

If our model is correct, the same principles should also apply to eukaryotic genomes. Indeed, eukaryotes also have mosaic genomes with a distribution of mutation rates, the main difference being that horizontal transfer in eukaryotes is much rarer than in prokaryotes [48]. We thus applied our method to the comparison of a few vertebrate genomes, and found that the resulting MLDs could be fitted with only the vertical part, see Fig. S9 (similar results were found in Refs. [38, 39, 49, 50]).

### Estimated parameters

Our model makes several predictions regarding the estimated parameters and their relationships. If the assumptions of our model are correct, we should observe these relationships as well in the empirical data. First, our model predicts that the total length of the homologous regions of the genomes of two species depends on the time divergence, see Eq. (6). For small time divergences, the full genome can be aligned, and after a certain time threshold (i.e. *τ > δ/μ*_*s*_), *L*_*v*_ decreases with the divergence time as *L*_*v*_*≃δ*^2^/(*μ*_*s*_*τ*)^2^, as predicted in Eq. (6). Indeed, this relationship is well reproduced on empirical data as shown in Fig. 3(a). Our model further predicts that the average divergence between the two genomes depends on the time divergence in a non trivial fashion, see Eq. (7). As predicted, we observe on empirical data that the average genome-wide divergence scales linearly for closely related species, reaches 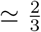 of the aligner detection limit *δ*for *τ* = *μ*_*s*_*/δ*, and then grows very slowly with *τ* (see Fig. 3(b)). Finally, we find that the rate of horizontal transfer, *ρ*, as presented in Fig. 3(c), can vary by orders of magnitude (see also [51]), and exhibits a clear trend to decay as the divergence time grows, as previously observed (see e.g. [44, 52–54]).

**FIG. 3:**
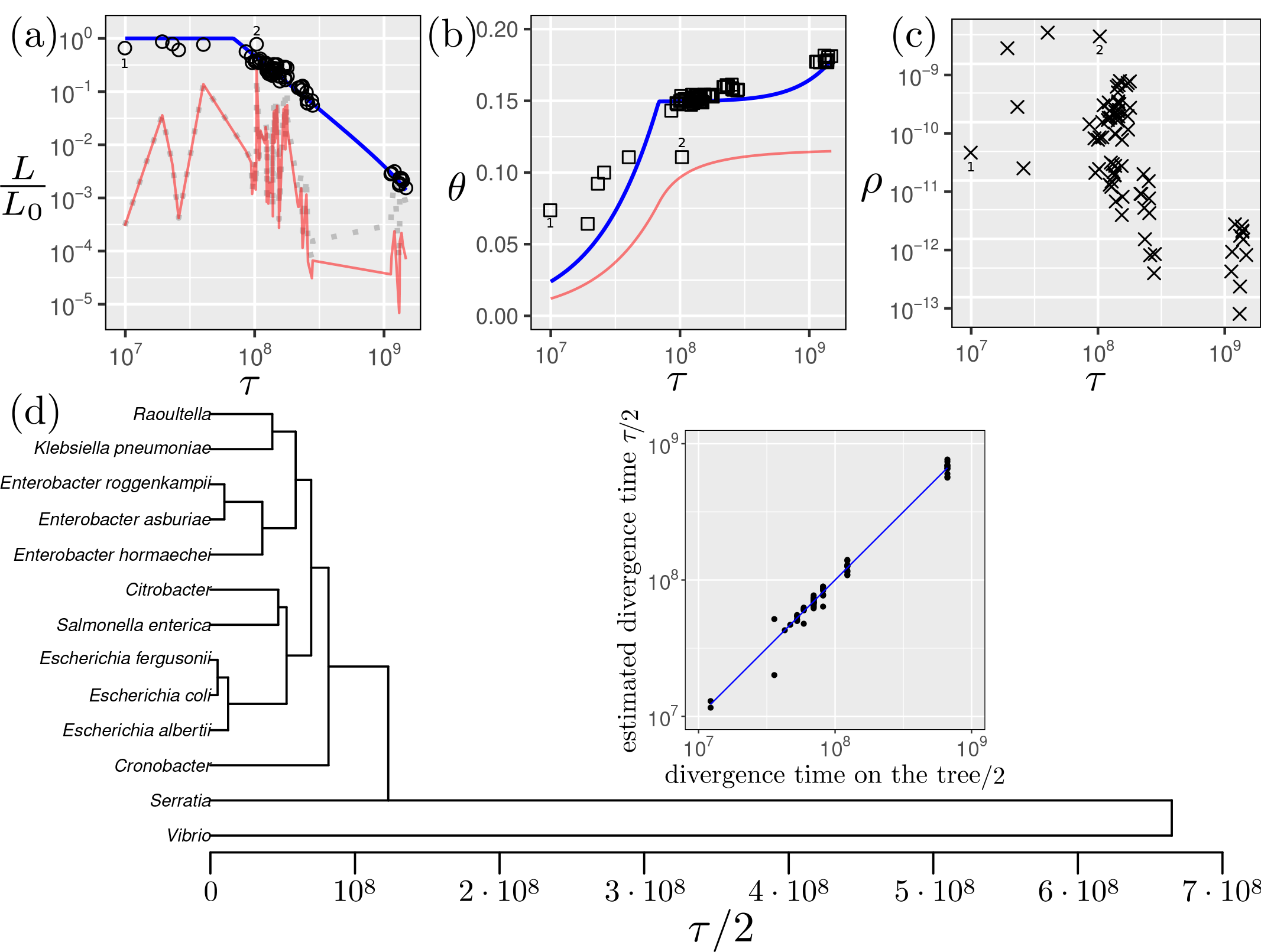
Relationships between the models parameters and the estimated time divergences for species of the *Enterobacteriaceae* family. (a) Ratio of the detectable homologous length and estimated value of the genome length of the common ancestor for all pairs of taxa as a function of the time divergence between the taxa. The blue line is *L*_*v*_*/L*_0_, the predicted length ratio from the vertical part based on Eq. (6), the red line is *L*_*h*_*/L*_0_ the predicted length ratio from the horizontally transferred and detectable part based on Eq. (9). The dotted grey line represents the full (detectable and non-detectable) length ratio of the horizontally transferred part, given by *ρτ* . The length ratio *L/L*_0_ from both detectable parts (vertically and horizontally transferred), from Eq. (12) is indistinguishable from *L*_*v*_—the blue line—on this scale for these data. Detailed empirical data for each taxa pair is shown in Fig. S5. (b) Empirical divergences for all taxa pairs after Jukes-Cantor distance correction [55] *vs*. fitted time divergence (squares). The blue line represents the predicted divergence along the vertical part (Eq. (7)), while the red line represents the predicted divergence along the horizontally transferred part (Eq. (10)). The total predicted divergence given by Eq. (13) is indistinguishable from *θ*_*v*_ for these data. Detailed empirical data for each taxa pair is shown in Fig. S6.(c) Fitted HGT rate as a function of the fitted divergence time for all pairs of taxa. Digits 1 and 2 on the upper panels indicate “model setbacks” for 1. *E. coli vs. E. fergusonii* and 2. *E. asburiae vs. E. hormaechei*. (d) UPGMA tree using estimated pairwise time distances *τ* . Inset plot compares the time distances on the resulting ultrametric tree with the estimated values (on a double-logarithmic scale).

Our model failed to estimate the time divergence for two specific taxa pairs: 1. *E. coli vs. E. fergusonii* and 2.*E. asburiae vs. E. hormaechei* (annotated with digits 1 and 2 in Fig. 3). In Fig. S10 one can see the reason for this: the MLD of the first pair has a *m*(*r*)*∼r*^*−*3^ regime in the vertical part, implying that Eq. (1) is not valid and suggesting instead *p*(*μ*)*∼μ*^0^ for this pair. One can observe similar behaviour for another closely related pair in Fig. 2(a). For the other pair, *E. asburiae vs. E. hormaechei* (see Fig. S10(b)), the rate of horizontal transfer is that high that the horizontally transferred loci dominate over the vertical part in the MLD. Since time divergence estimation is based solely on the vertical part, the signal is obfuscated. In fact, this demonstrates another quality of our approach: analysing the MLD, one can easily diagnose special pairs of taxa for which estimating the parameters requires a different approach. Overall, the empirical data agree well with our theoretical predictions, demonstrating the accuracy of the model.

### Resulting phylogenetic tree

We next investigated whether the time estimates we computed with our method were coherent and compatible. To do so, we computed a phylogenetic tree from our time estimates using the UPGMA method [56], forcing ultrametricity of the tree (see Methods). The resulting phylogenetic tree is shown in Fig. 3(d). In the inset of the figure one can see that the pairwise distances in the obtained tree follow closely the estimated values. This result shows that our estimated time divergences have an inherent ultrametric structure, demonstrating that all considered lineages have similar mutation rate distributions. Moreover, the topology of the tree reproduces well what is expected based on the literature: for instance, *Klebsiella pneumoniae* is closely related to *Raoultela* [57] and *S. enterica* is closely related to *Citrobacter* [58].

In contrast, using a simple molecular clock assumption, the topology of the tree and the value of the time estimates would be very different (see Fig. S4). Indeed, because segments with high divergence are not identified by the alignment software, the estimated divergence are greatly underestimated for distant pairs, and the resulting tree would have an unresolved star-like structure.

## IV. Discussion and summary

In this paper, we studied the statistical properties of similarities between bacterial genomes. Similarities between two bacterial genomes are shaped by *mutations, horizontal transfer of genes/alleles, gene losses* and *selection* during their evolution since their last common ancestor. In practice, the observed similarities are also shaped by the *sensitivity* of the used aligner: if two loci are too evolutionary distant, the aligner cannot detect their homology and these distant loci are disregarded. In this case, only the more conserved loci are detected by the aligner, making the bacterial genomes appear more similar than they really are.

In this study we combine all these factors and propose a mathematical framework to model and assess their contributions. We show that the analysis of match length distributions is a powerful tool that reflects details of bacterial evolution. In our model, *mutations* are assumed to occur randomly, breaking long matches to shorter ones.Different loci mutate with different effective rates, *μ*, depending on their associated *selective* pressure. On the other hand, *horizontal transfers* between two genomes generate long matches with a given rate *ρ. Gene losses* reduce the total length of homologous loci, scaling down the MLD prefactor *L*_0_. The *sensitivity* of the aligner is modelled by considering only homologous loci with an average divergence lower than a threshold *δ*.

We find that the shape of the mutation rate distribution strongly influences the size of the alignable part of the genomes, and, as such, the average divergence between them. By explicitly modeling the distribution of mutation rates along genomes, we can resolve the long-standing discrepancy between the spontaneous mutation rate measured in short time-scale experiments and the one inferred from distant bacteria on the evolutionary time-scales without the *ad hoc* assumption of time-dependent mutation rate. Our results indicate that the mutation rate distribution is linear (see Fig. 1(b)) between two extreme values: the spontaneous mutation rate *μ*_*s*_ and the mutation rate of the most conserved loci *μ*_*c*_.

### Selective pressure and distribution of effective mutation rates

The distribution of mutation rate along genomes reflects the variation of selective pressure. The selective pressure on a locus is affected by the fitness effect of a mutation at this locus and by the effective population size of the taxon [59, 60]. The distribution of fitness effects can in principle be assessed [16], but these methods require in general to conduct complex experiments in controlled environments. In contrast, in this study we directly model the mosaic distribution of effective mutation rates under simple assumptions.

If mutation rates along different lineages are not correlated, one expects that the mean effective mutation rates are linearly distributed (see Eq. (1)) [38, 39]. The validity of the no-correlation assumption is not obvious—more conserved loci in one lineage are expected to tend to be more conserved in another lineage. Therefore, there might be another explanation for the approximate linear distribution of the effective mutation rates that we observe empirically in very different comparisons: from bacteria to mammals (see Fig. S9). This aspect requires more detailed study, for instance analysing bacterial ancient DNA [1].

### HGT detection based on MLD

Detection of horizontal transfer of a locus is often based on its high similarity in two organisms, much higher than one would expect due to conservation [61]. Long (almost) exact matches are often interpreted as horizontal transfers, [44, 53, 62, 63]. However, in the absence of a model, it is not clear what is the threshold that discriminates horizontally transferred and well conserved sequences, leading to false-negative or false-positive detection errors [64]. Detailed analysis of the MLD can help to minimize those errors: the presence of two clear regimes in the MLD suggests that sequences with exact matches shorter than the crossover between the two regimes are conserved, while longer ones most probably have been horizontally transferred. This way of classifying loci is non-parametric and can be applied to almost all pairs of taxa studied here (see below for exceptions), which all exhibit MLDs with two clear regimes.

Since the model proposed here clearly disentangles conservation and horizontal transfers, our estimate of horizontal transfer rates are expected to be more accurate than the one found in previous studies. Our results confirms that the rate decreases with the divergence time, as previously observed [44, 51–54]. Note, that in this study we filtered out plasmid sequences, so that the estimated horizontal transfer rates are related only to the chromosomal part of the genome.

### Phylogenetic analysis

Using the presented approach we built an ultrametric tree of the *Enterobacteriaceae* family with *Serratia* and *Vibrio* as outgroups (see Fig. 3(d)). Topologically the tree reproduces known phylogenetic relationships.

While these relationships can also be found on a tree constructed using the average genome-wide divergences (see Fig. S4), we emphasise that this tree is not topologically identical to the one built using our method. For instance, the average divergence tree suggests that *Salmonella* is closer to *Klebsiella* (*θ* = 0.1479) than to *Escherichia* (*θ* = 0.1498).The tree based on the time divergences estimated using our method suggests the opposite (*τ* = 1.88·10^8^ and *τ* = 6.67·10^7^, respectively), in agreement with the 16S rRNA result [65, 66] (although 16S rRNA phylogeny cannot be taken as a ground truth [27, 67]).

Overall, for closely related taxa where the molecular clock still holds, the two methods yield very similar trees. In contrast, for distantly related taxa where many homologous sequences are too diverged to be identified by the aligner, the branch lengths estimated by our method are very different from those found by the average divergence method. As a consequence, the average divergence tree has a star-like shape, while our method can better resolve deep branching patterns.

Interestingly, we estimate that *Escherichia* and *Salmonella* branched 100·10^6^yrs ago. This is ≃30% earlier than the currently accepted estimate of 140·10^6^yrs based on the appearance of mammals [29]. This suggest that our assumptions about the values of the mutation rates *μ*_*c*_ and *μ*_*s*_ are higher than the real ones or that the branching of the two taxa occurred significantly after the appearance of mammals.

### Model Setbacks

For two taxa pairs the time divergence estimates are not accurate. Possible reasons for this may be that the recombination rate between the two species is so high that the vast majority of the observed matches result from horizontally transferred loci rather than from evolutionary conserved ones. Hence, one cannot reliably estimate the time divergence using our approach. Another possible reason might be that the mutation rate distribution does not follow the linear distribution assumed in our model. Although the assumption of linear mutation rate distribution is very general and is fulfilled in most cases, it might be violated for closely related pairs in at least two scenarios: *(i)* if the effective mutation rates of the homologous loci are well correlated or *(ii)* if mutations occur mostly at loci with high effective mutation rates for which the asymptotic scaling considerations in Section II do not apply.

### Improvement of the method

For simplicity, in this article, we used a strict molecular clock, meaning that the mutation rate distribution is the same along all branches, and we assumed that the mutation rate varies only along the genome. We demonstrate that this simple model is consistent with the current knowledge of bacterial evolution and allows to estimate reasonable divergence times. The presented approach could further be extended, relaxing the clock also along the lineages by assuming a different distribution of the effective mutation rate along every branch. This extension has the potential to improve the quality of the parameter estimations, and solve some discrepancies of our model (see discussion above), although it would require to make many more *ad-hoc* assumptions and to fit a much larger number of free parameters.

On the technical side, in this paper, we used the `nucmer` software to construct all alignments, because this method is computationally very efficient. However, our framework could easily be adapted to other algorithms with improved sensitivity (*e*.*g*. `lastz` [68]) to align more distantly related genomes and measure horizontal transfer rates and time divergences. The exact same theoretical framework could be used, just changing the effective parameter *δ*to account for the difference in sensitivity.

### Summary

We demonstrated that a method embracing the complex mosaic structure of bacterial genomes and explicitly accounting for the technical limitations of sequence homology detection can improve the estimation of deep phylogenetic branches and their timing. The main advantage of our method is that it can leverage genome-wide alignment data resulting in robust time divergence estimates that are not dependent on a few specific marker genes. Our results have implications that go beyond bacterial evolution as we have shown that our model applies to the mosaic structure of many more genomes, including vertebrates.

## V. Methods

Throughout this article the time (*τ*) units are years, length (*L*) units are bp, while the rates (mutation *μ* and horizontal transfer *ρ*) are in units of yrs^*−*1^ bp^*−*1^.

### Data

To validate the theoretical predictions we used taxa from the well known *Enterobacteriaceae* family and two outgroups: *Serratia* and *Vibrio* genera. We considered only chromosomal part of the genome. To filter out plasmids we used only complete genome and chromosome level assemblies in the RefSeq [69] and GenBank [70] databases using NCBI [71]. We considered only species with at least 20 available assemblies. For species with smaller number of full assemblies we grouped the species to corresponding genera.

### Aligning the genomes

To align pairs of genomes we used `nucmer` [47] with the default settings, using only unique matches in both genomes (`--mum` option). To estimate the divergence we calculated the number of differences per alignment, normalized by the alignment length. We consider all insertions and deletions as a single difference. The code is available on the `github` repository [72].

### Plotting MLDs – log binning

To plot the MLD we used a linear 3bp binning up to 35bp and logrithmic binning with 10 points per decade for larger matches. Namely, our breaks of the histogram are: 0.5, 2.5, 5.5, 8.5, …, 35.5, 35.5·10^0.1^, 35.5·10^0.2^, 35.5·10^0.3^… Within each bin we count the number of matches and normalize it by the size the bin. In addition, we normalize the MLD by the total number of alignments we do for the two considered taxa. If we analyze two taxa with *n*_1_ and *n*_2_ genomes, respectively, we do *n*_1_*×n*_2_/2 alignments, collect all the exact matches and, therefore, divide the total MLD by *n*_1_*×n*_2_/2. The code is available on the `github` repository [72].

### Simulating the genomes

To simulate bacterial evolution we started from 5,005,213-long *E. coli* chromosome NZ_CP092647.1 and divided it to segments with different lengths, distributed exponentially with an average of 10^4^bp. Each segment was evolved with a mutation rate drawn from the distribution of Eq. (1) with *μ*_*c*_ = 10^*−*10^ and *μ*_*s*_ = 3.64·10^*−*9^. We assume that transversions and transitions occur with the same probability and back mutations are allowed. We used this framework to evolve pairs of genomes with a wide range of divergence times, from 10^7^yrs to 9·10^8^yrs. Horizontal transfer is implemented by transferring a random segment of length 10^4^ from one branch to another with rate *ρ*= 10^6^yrs*/τ* ^2^ per bp, to mimic the relationship between the horizontal transfer rate and the divergence time observed in real data. For each value of *τ* we simulated 12,800 pairs with different random seeds. The resulting sequences were aligned and analyzed using the same procedure used for empirical genomes (see Sections V and V). The code is available on the `github` repository [72].

### Segmenting the genomes (segmut package)

Our model assumes mosaic structure of the genome: different loci mutate with different rate and may have different divergence time due to horizontal transfer. Our approach allows to analyze the evolutionary history without explicit identification of the mosaic segments. However, to get more confidence about our approach and the results, we segmented the alignments of the bacterial genomes with respect to the density of mutations. To do so we followed the ideas in Refs. [73–75], maximizing the *?*^2^ statistic of the mutations density of the segments. The R package implementation can be found in the github repository [41].

### Fitting procedure

To fit the empirical MLD using Eq. (11) for each pair of taxa, we used two free parameters: *τ* and *ρ*. Genome length of the common ancestor of the pair of taxa, *L*_0_, is taken as the length of the smallest genome of the pair. The fitting is performed by minimizing the mean square relative difference between the theoretical and the empirical MLDs using the Nelder-Mead algorithm [76]. To find the best starting point we used Harmony Search heuristic [77] with 10,000 starting points. The code is available on the `github` repository [72].

### Building the tree

We build the ultrametric tree using hierarchical clustering of the taxa based on their estimated pairwise time divergences *τ* . We use average linkage clustering (`hclust` function from the `stats R` package), which is equivalent to the UPGMA method [56]. To get the pairwise distances from the resulting tree we use `cophenetic` function from the `stats R` package.

## VI. Acknowledgements

Authors thank M.S. Gelfand for useful comments and discussion. Numeric analysis was carried out using the supercomputer cluster “Afalina” in Sevastopol State University.

## A. Supplementary material

**FIG. S1:**
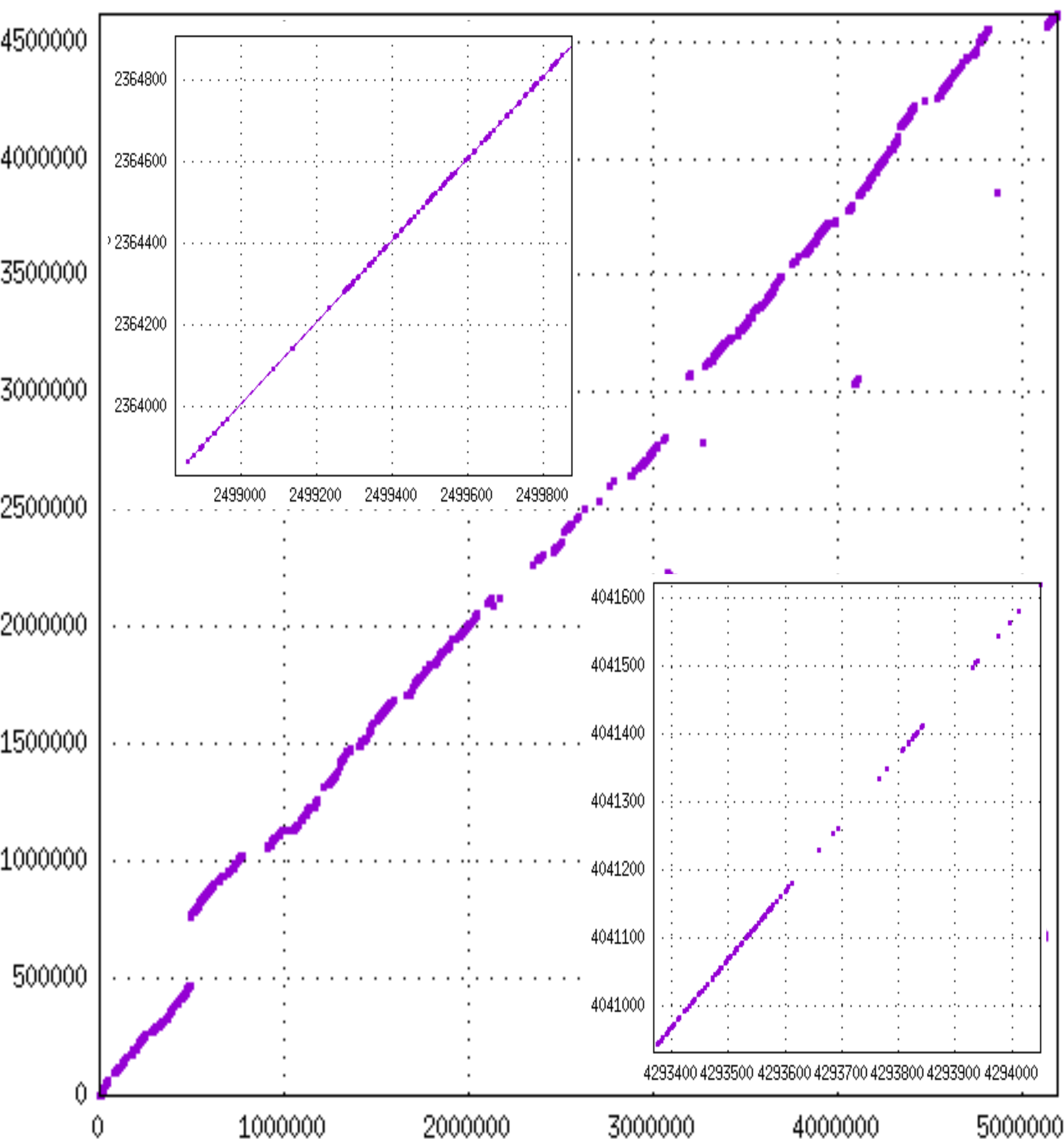
Dotplot using `nucmer` of *E. coli* (NZ_CP068796.1 strain RIVM_C012087) and *S. enterica* (NZ_CP019035.1 str. 9184 isolate ATCC 9184). The alignment was done using `nucmer` with the default settings and `--mum` option and visualized using `gnuplot` [78]. Markers denote differences between the two homologous genomic loci.

**FIG. S2:**
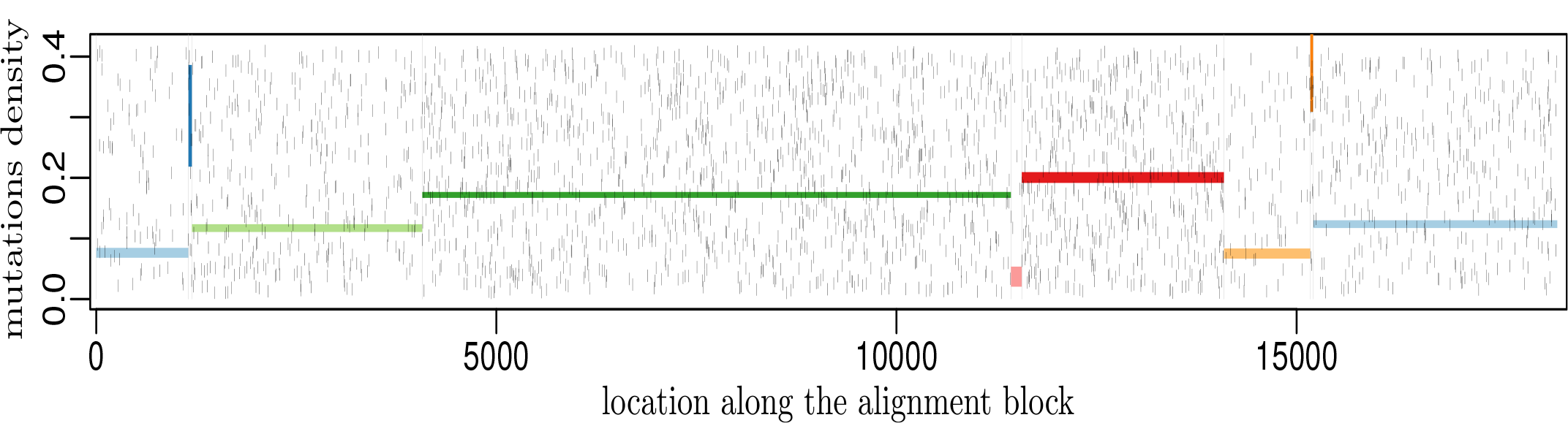
Mutations along the largest alignment block for *E. coli* strain ATCC 8739 (NZ_CP033020.1) and *S. enterica* strain SE20-72C-2 (NZ_AP026948.1). Each short vertical black line represents a mutation (vertical position is random). Colored rectangles represent the 68% confidence intervals of the mutation density along the segments (*±*one standard deviation, calculated assuming Poisson distribution), obtained using the `segmut R` package.[41]

**FIG. S3:**
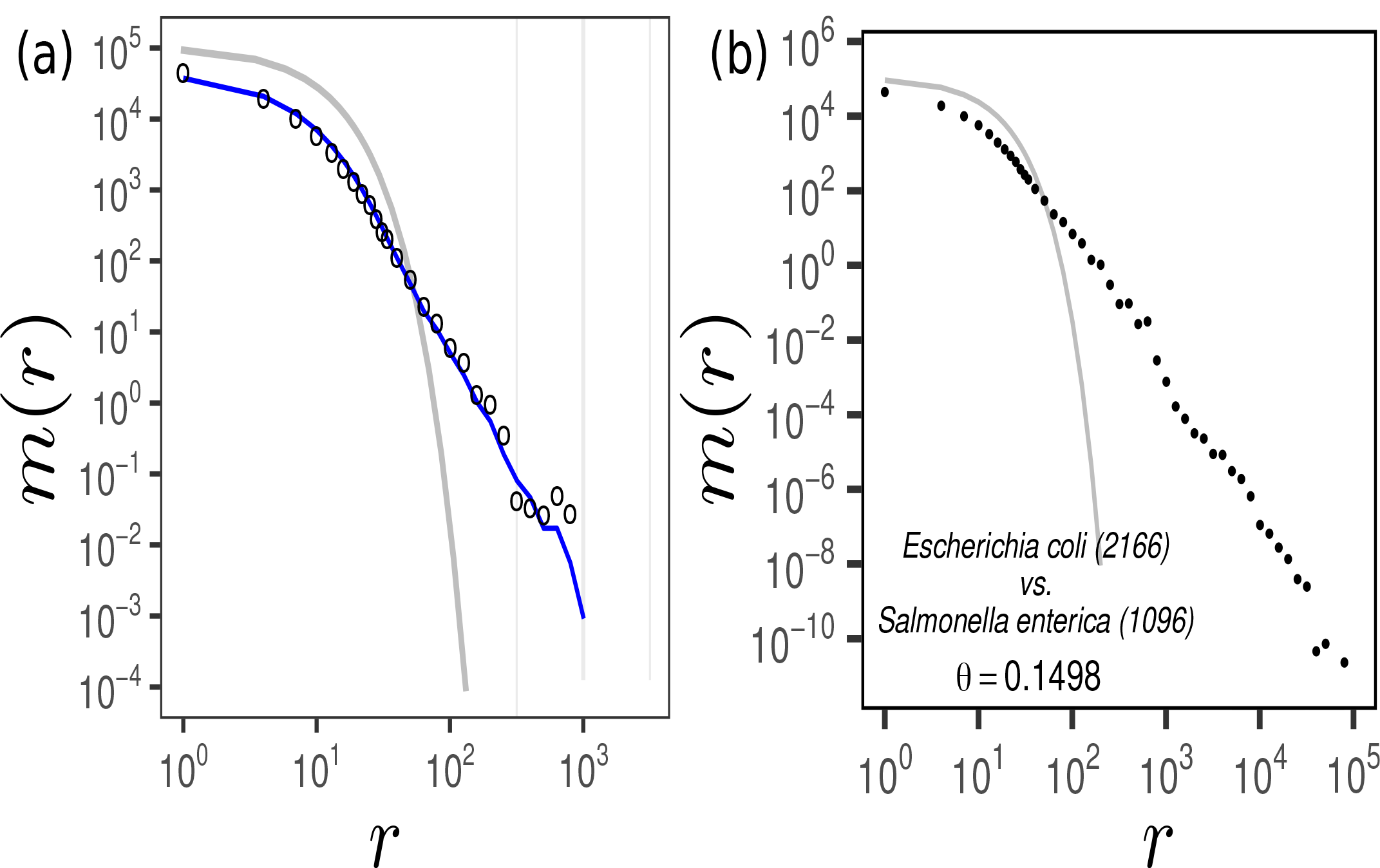
(a) Empirical *vs*. pseudotheoretical MLD. Circles represent empirical MLD for alignment of *E. coli* strain ATCC 8739 (NZ_CP033020.1) and *S. enterica* strain SE20-72C-2 (NZ_AP026948.1). The line is based on the segmentation of the alignment using the `segmut R` package. For each segment *i* the divergence *θ*_*i*_ and length *K*_*i*_ are calculated and then the pseudotheoretical MLD is calculated as 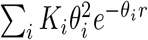 . (b) MLD calculated from all *vs*. all 2, 166×1, 096 alignments of *E. coli vs. S. entrerica* strains. Grey solid line in both panels represents theoretical MLD, ignoring mosaic structure of the genome, assuming that the average genome-wide density of mutations *θ* is uniform along the genome, *Lθ*^2^*e*^*−θr*^ (see Eq. (2)).

**FIG. S4:**
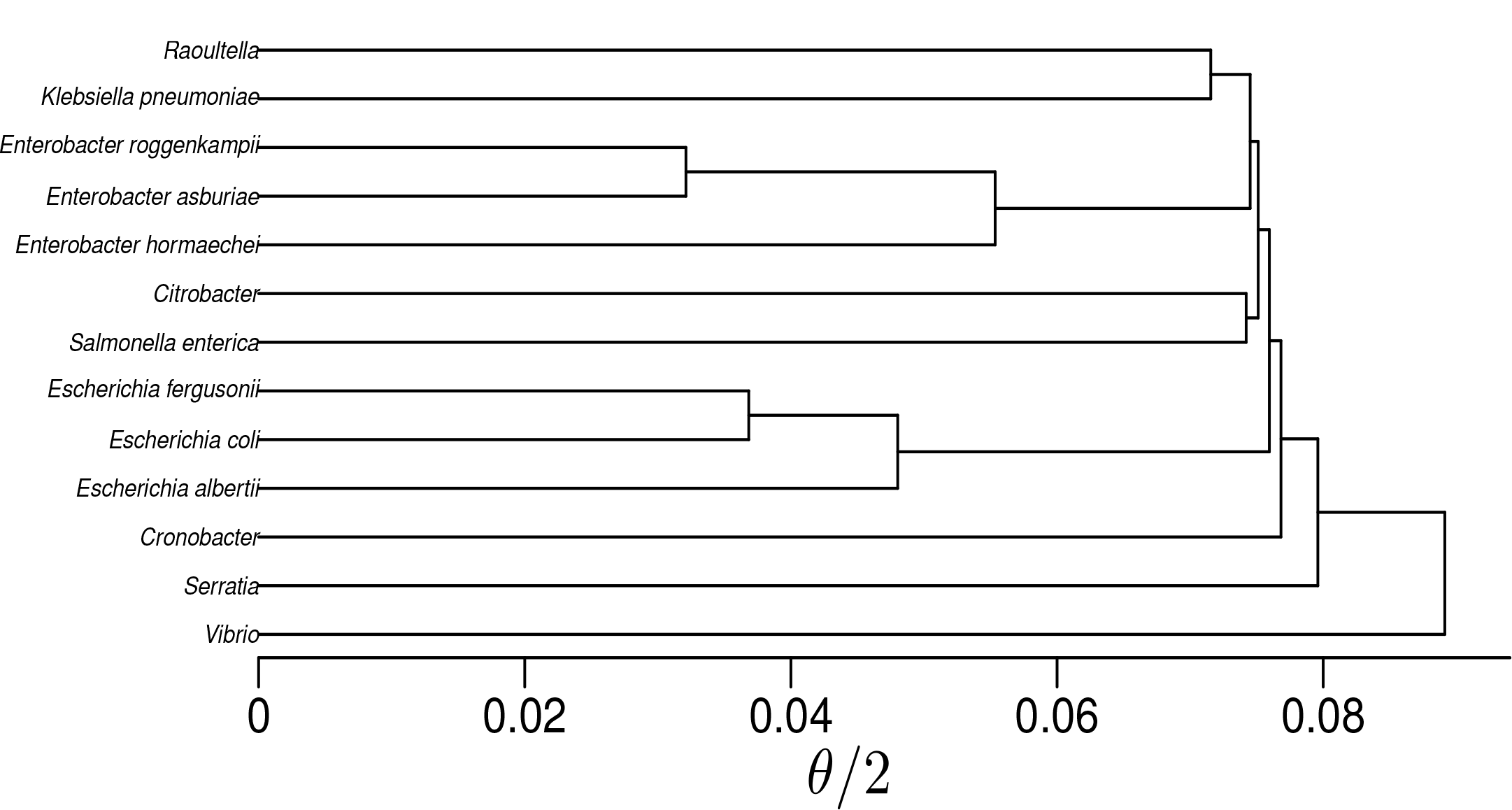
UPGMA tree using the average divergences *θ*.

**TABLE I:** Number of fully assembled chromosomal genomes used in this study for each taxon.

|  |  |
| --- | --- |
| *Escherichia coli* | 2,166 |
| *Klebsiella pneumoniae* | 1,226 |
| *Salmonella enterica* | 1,096 |
| *Vibrio* | 619 |
| *Enterobacter hormaechei* | 279 |
| *Citrobacter* | 192 |
| *Serratia* | 140 |
| *Escherichia fergusonii* | 50 |
| *Enterobacter roggenkampii* | 39 |
| *Raoultella* | 32 |
| *Enterobacter asburiae* | 25 |
| *Escherichia albertii* | 22 |
| *Cronobacter* | 22 |

**FIG. S5:**
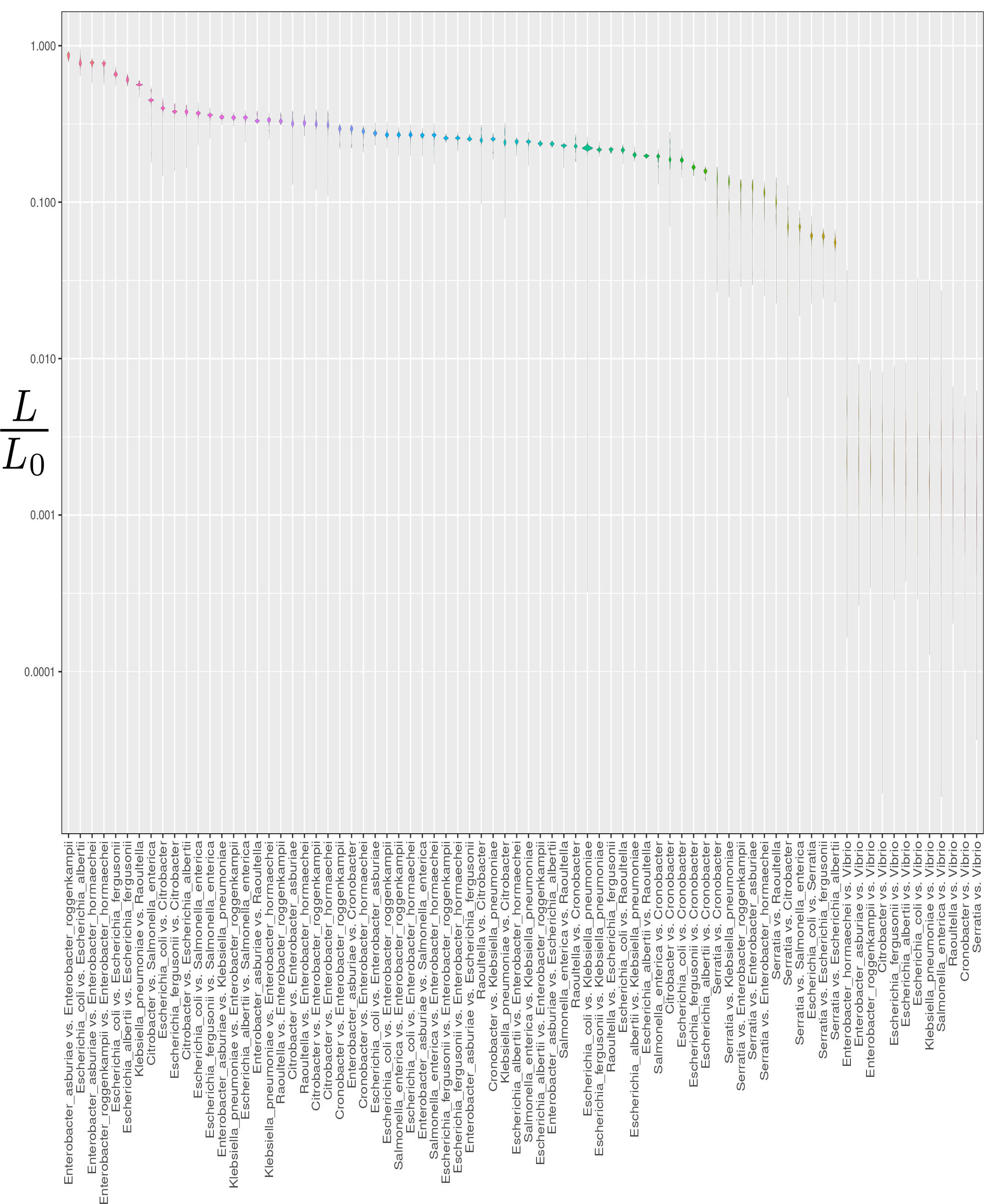
Violin plot of homologous fraction of the genome, detected by the aligner for all pairwise alignments ordered by the median value.

**FIG. S6:**
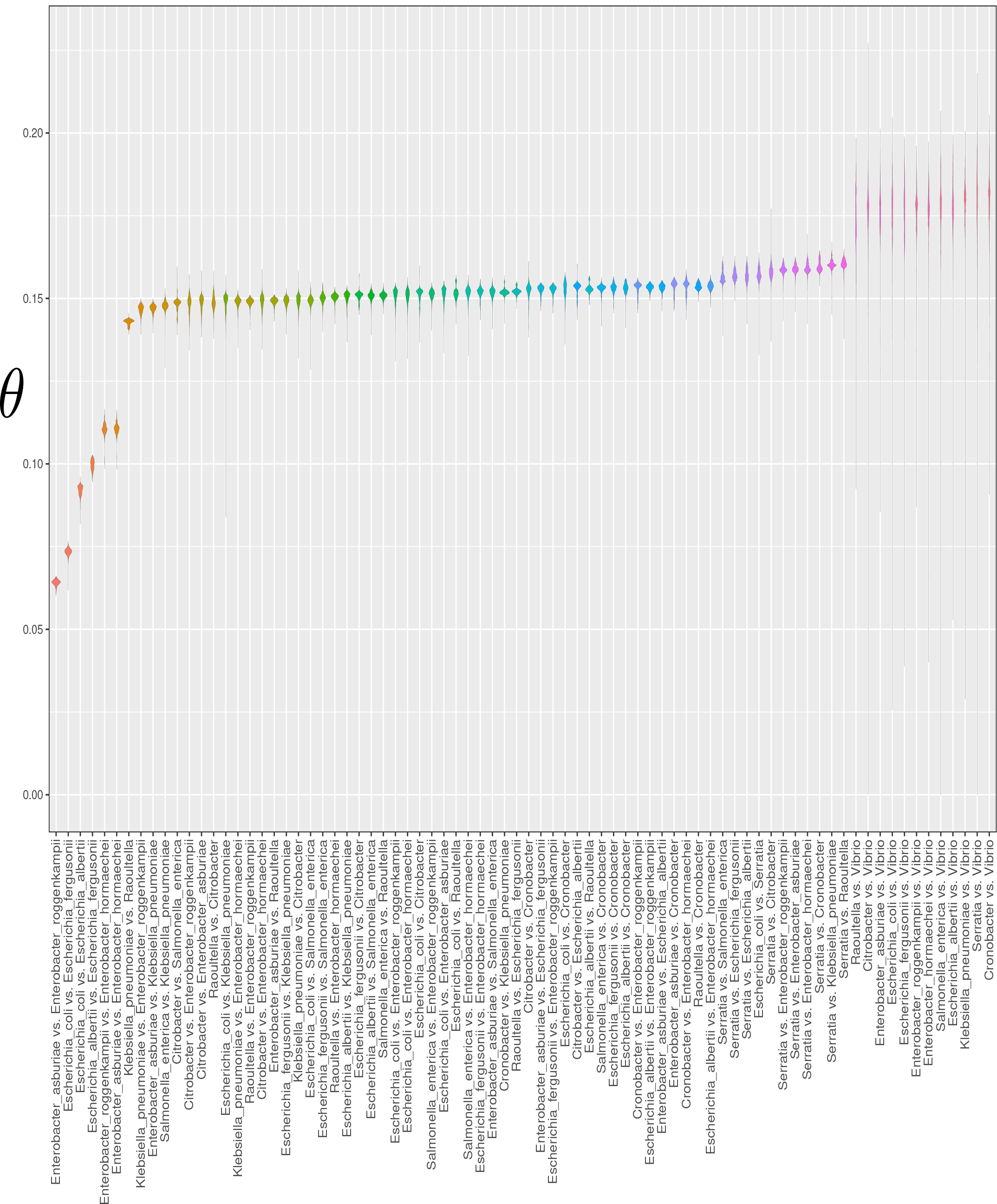
Violin plot of average divergences *θ* for all pairwise alignments ordered by the median value.

**FIG. S7:**
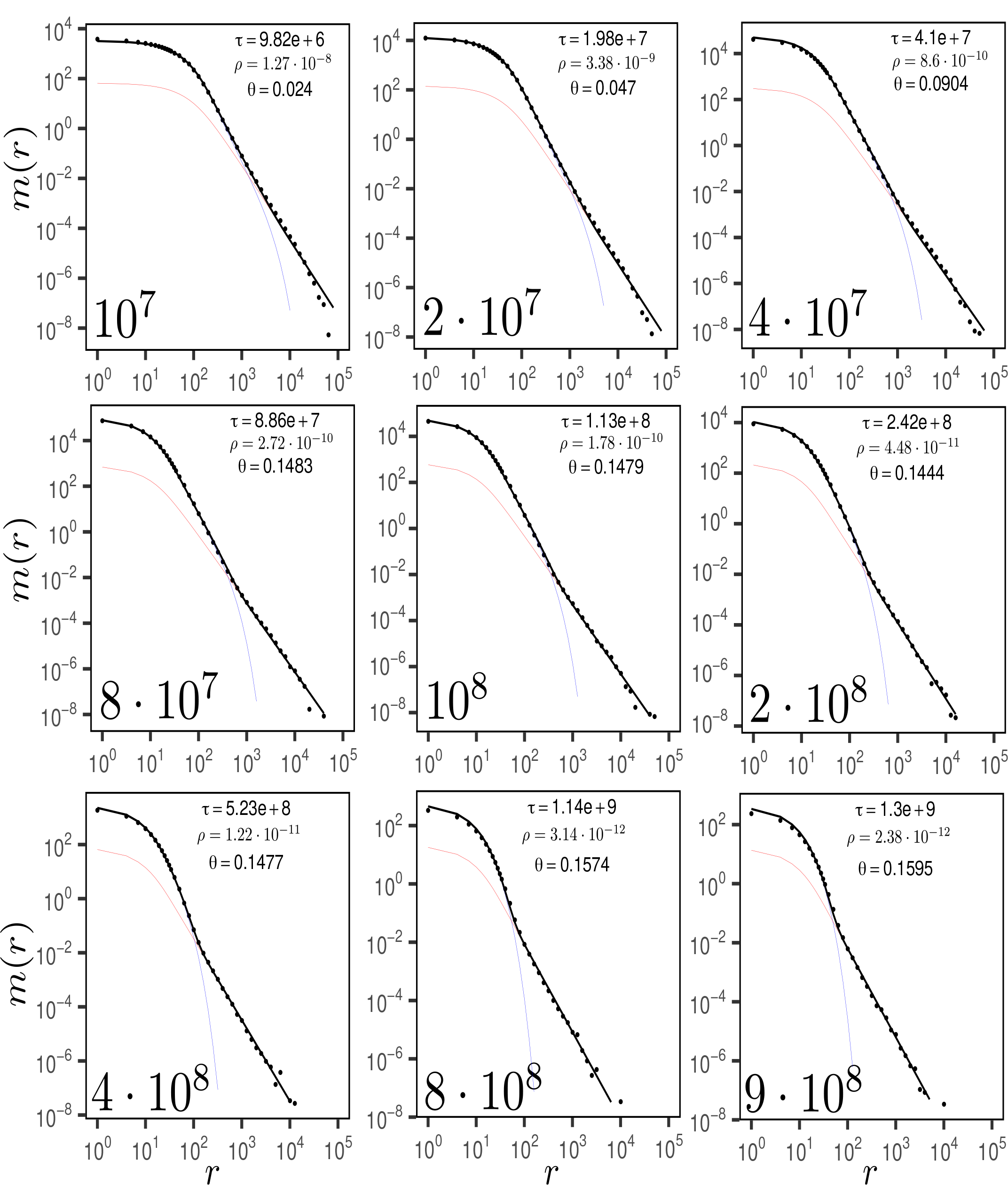
Match length distributions of simulated pairs of genomes with different divergence times, indicated in the bottom-left corner of the panels. The average divergence is indicated by *θ*. The simulated data (dots) are fitted using the global parameters *μ*_*c*_ = 10^*−*10^ and *μ*_*s*_ = 3.64·10^*−*9^ and *δ*= 0.25. The values of *τ* and *ρ*are fitted for each pair separately and are shown in the top-right corner.

**FIG. S8:**
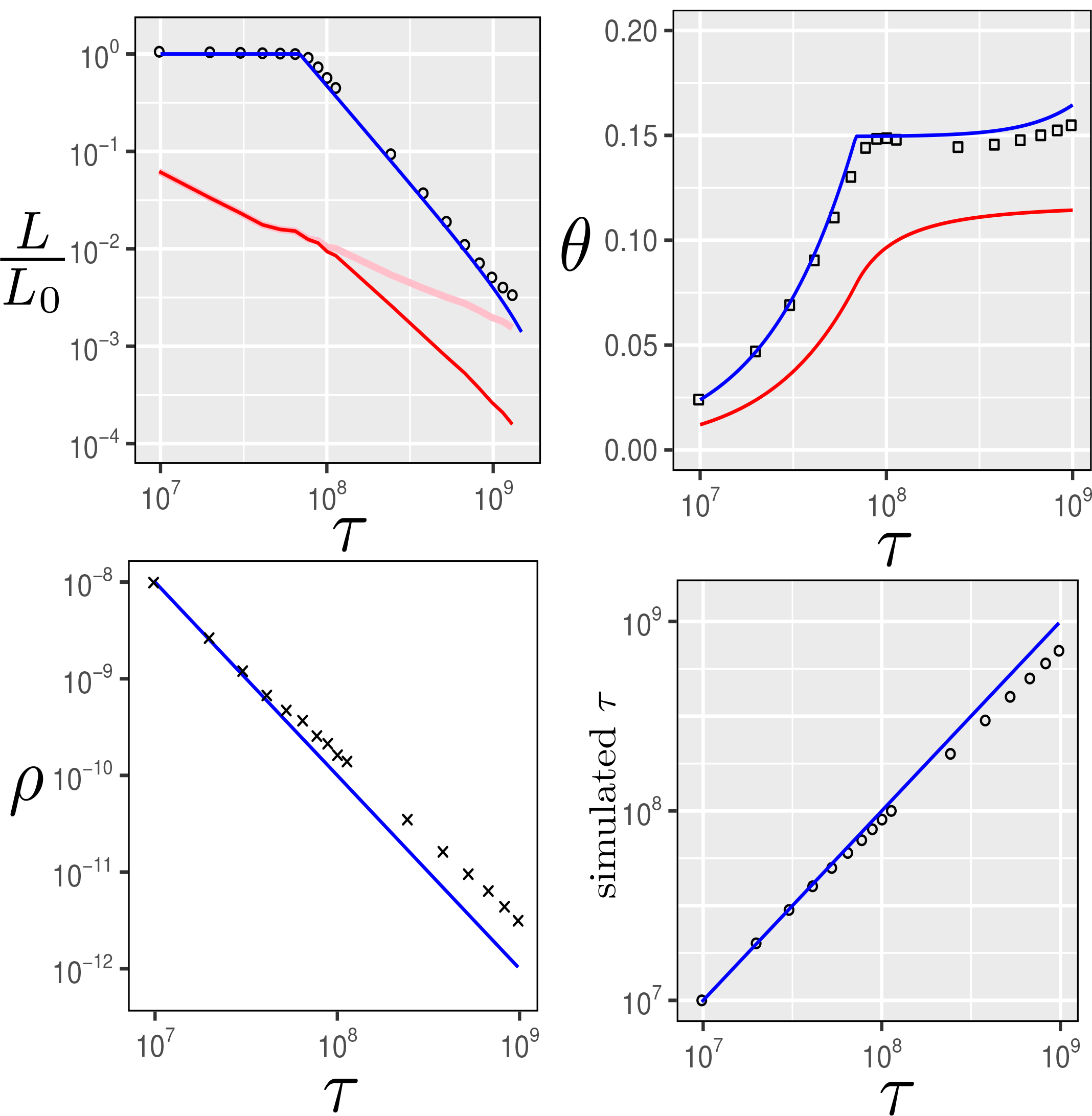
Analysis of the simulated sequences. (a) Ratio of detectable homologous length and fitted value of the genome length of the common ancestor for all pairs of taxa as a function of fitted time divergence between the taxa. Blue line is *L*_*v*_*/L*_0_, the predicted length ratio from the vertical part based on Eq. (6), red line is *L*_*h*_*/L*_0_ the predicted length ratio from the horizontally transferred and detectable part based on Eq. (9). Pink line is the full (detectable and non-detectable) length ratio from the horizontally transferred part: *ρτ* . The length ratio *L/L*_0_ from both detectable parts (vertically and horizontally transferred), from Eq. (12) is indistinguishable from *L*_*v*_—the blue line—on this scale for these data. (b) Empirical divergences for all taxa pairs after Jukes and Cantor distance correction [55] *vs*. fitted time divergence are indicated by squares. Blue line represents the predicted by Eq. (7) divergence along the vertical part, while the red line represents the predicted by Eq. (10) divergence along the horizontally transferred part. The total predicted divergence given by Eq. (13) is indistinguishable from *θ*_*v*_ for these data. (c) Fitted HGT rate as a function of the fitted divergence time for all pairs of taxa.

**FIG. S9:**
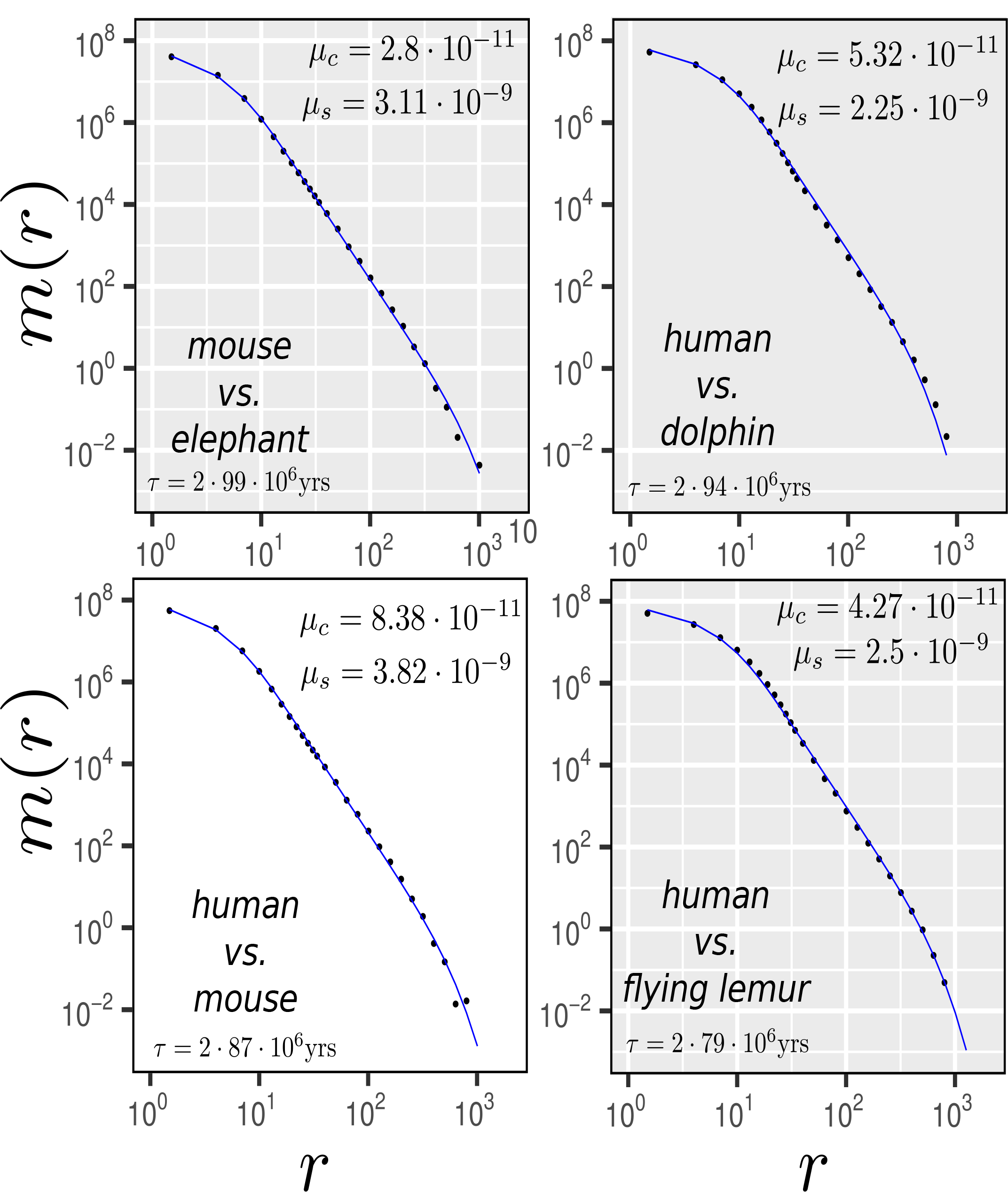
MLD of animal pairs alignments, downloaded from UCSC [79] (note that here the alignment was done not by `nucmer`, but using chained `lastz` aligner [68, 80]). The analytical fit of *μ*_*c*_ and *μ*_*s*_ (see upper-right corner) was done using (5)—the vertical part of the MLD (we assume that here there is no horizontal transfer). The divergence times *τ* are taken from Ref. [81] (see bottom-left corners).

**FIG. S10:**
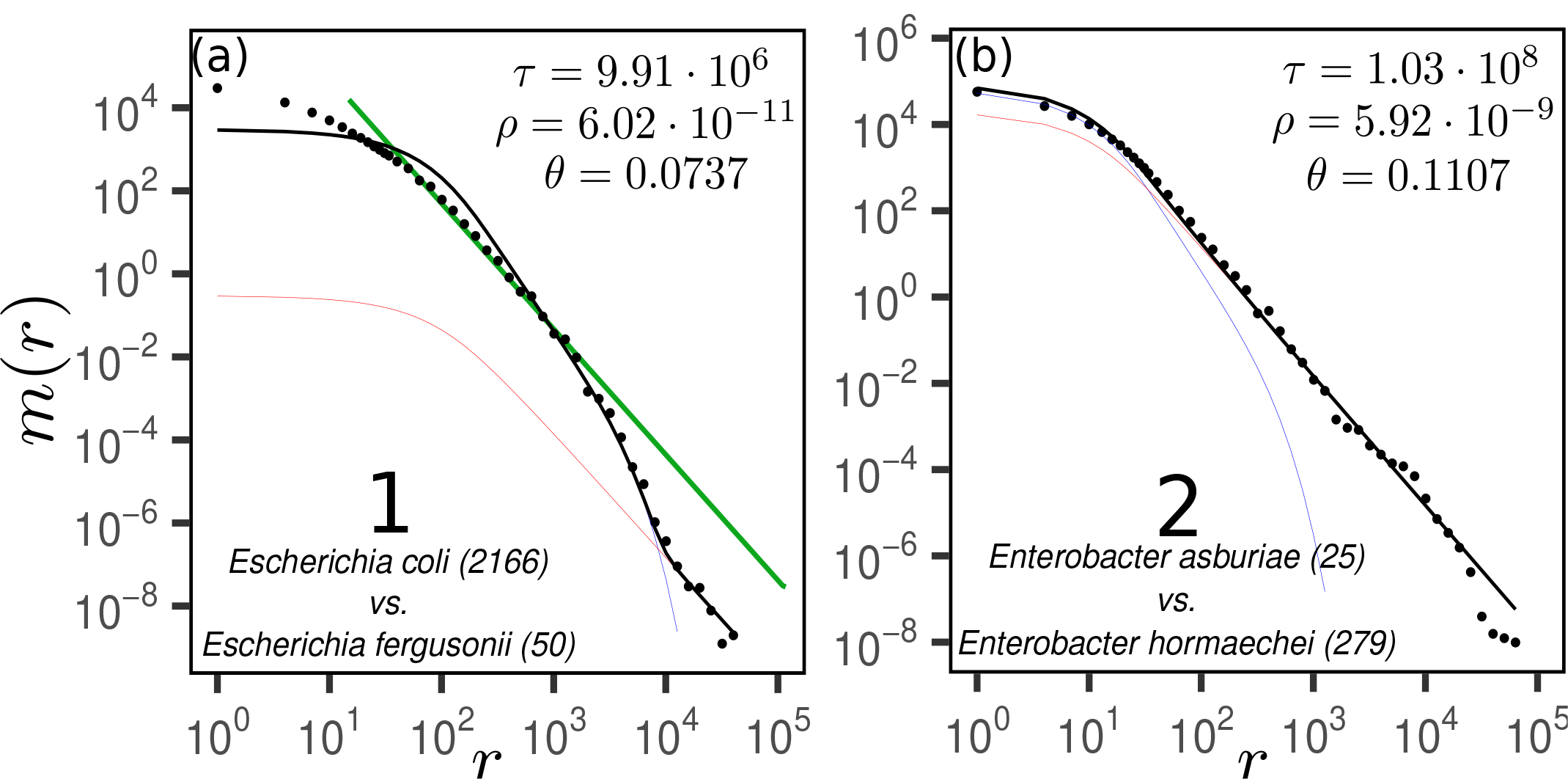
Match length distributions of 2 pairs of taxa for which the model assumptions are not valid and the time divergence estimation is not accurate. Names of the taxa are indicated in the bottom-left corner of both panels. The numbers in the brackets indicate number of strains. The average divergence is indicated by *θ*. The empirical data (dots) are fitted with Eqs. (5,8,11) using the global parameters *μ*_*s*_ = 3.64·10^*−*9^/bp/yrs, *μ*_*c*_ = 10^*−*10^/bp/yrs and *δ*= 0.25. The values of *τ* and *ρ*are fitted for each pair separately and are shown in the top-right corner. The genome length of the most recent common ancestor of two taxa is assumed to be the minimum of the taxas’ genome lengths. The green line in (a) corresponds to *m*(*r*) *∼ r*^*−*3^ power-law.

**FIG. S11:**
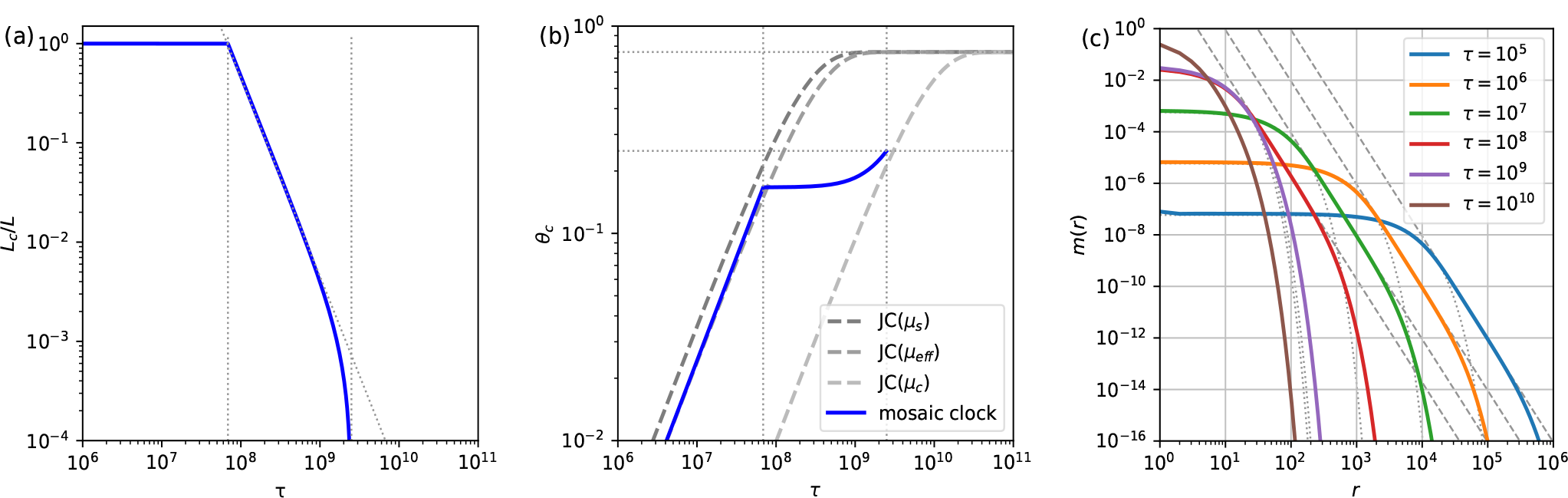
(a) the length of the *δ*-detectable fraction of a genome. The vertical lines are at *δ/μ*_*s*_ and *δ/μ*_*c*_. The detectable part decreases proportional to 1*/τ* ^2^ initially as indicated by the dotted line. (b) The empirical divergence as a function of *τ* . The dashed lines represent the prediction due to the Jukes-Cantor model with rates 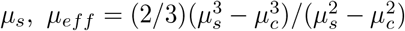, and *μ*_*c*_. (c) Match length distributions for several values of *τ* . The functions for small *τ < δ/μ*_*s*_ show a power-law regime with exponent *−*4, as indicated by the dashed gray lines. Corresponding exponential distributions with the same mean are shown with gray dotted lines.

